# Tandem tyrosine residues in the EPEC multicargo chaperone CesT support differential type III effector translocation and early host colonization

**DOI:** 10.1101/270066

**Authors:** Cameron Runte, Umang Jain, Landon J. Getz, Sabrina Secord, Asaomi Kuwae, Akio Abe, Jason J. LeBlanc, Andrew W. Stadnyk, James B. Kaper, Anne-Marie Hansen, Nikhil A. Thomas

## Abstract

Enteropathogenic *Escherichia coli* (EPEC) are worldwide human enteric pathogens inflicting significant morbidity and causing large economic losses. A type 3 secretion system (T3SS) is critical for EPEC intestinal colonization, and injection of effectors into host cells contributes to cellular subversion and innate immune evasion. Here, we demonstrate that two strictly conserved C-terminal tyrosine residues, Y152 and Y153, within the multicargo T3SS chaperone CesT serve differential roles in regulating effector secretion in EPEC. Conservative substitution of both tyrosine residues to phenylalanine attenuated EPEC type 3 effector injection into host cells, and significantly limited Tir effector mediated intimate adherence, a key feature of attaching and effacing pathogenesis. Whereas CesT Y153 supported normal levels of Tir translocation, CesT Y152 was strictly required for the effector NleA to be expressed and subsequently translocated into host cells during infection. Other effectors were observed to be dependent on CesT Y152 for maximal translocation efficiency. Unexpectedly, EPEC expressing a CesT Y152, Y153F variant exhibited significantly enhanced effector translocation of many CesT-interacting effectors, further implicating Y152 in CesT functionality. A mouse infection model of EPEC intestinal disease using *Citrobacter rodentium* revealed that CesT tyrosine substitution variants displayed delayed colonization and were more rapidly cleared from the intestine. These data demonstrate genetically separable functions for strictly conserved tandem tyrosine residues within CesT. Tyrosine 152 of CesT is implicated in NleA expression, providing functional relevance for localized amino acid conservation. Therefore, CesT via its novel C-terminal domain, has relevant roles beyond typical T3SC that interact and stabilize effector proteins.

## Importance

Pathogens with type III secretion systems absolutely require dedicated chaperone proteins to mediate efficient effector protein injection into host cells during infection. While these chaperones are well established as effector binding partners in the bacterial cytoplasm, their roles in regulating effector protein injection are modestly characterized. In the case of multicargo chaperones that bind multiple effectors, it is unknown how effector trafficking is coordinated during infection. In this study, we implicate two strictly conserved C-terminal tandem tyrosine residues of CesT in differentially regulating effector secretion in EPEC. Moreover, we discovered that tyrosine 152 of CesT was required for the expression of an effector protein. These findings further expand the importance of CesT in context of EPEC pathogenesis, revealing an unexpected site-specific functionality for a type III secretion chaperone.

## Introduction

Enterohaemorrhagic and Enteropathogenic *Escherichia coli* (EHEC and EPEC) are worldwide diarrheal pathogens that are associated with significant morbidity, and in extreme cases mortality (1–4). The <u>T</u>ype <u>3</u> <u>S</u>ecretion <u>S</u>ystem (T3SS) of these and other attaching and effacing (A/E) pathogens is a critical feature for host intestinal colonization (5–8). These bacteria use their respective T3SS to inject a variety of effector proteins directly into host cells thereby subverting host cell functions and enabling the bacteria to evade the host’s innate immune responses (9). A hallmark of A/E pathogen gut infection is microcolony formation and intestinal lesions. Furthermore, strains that express Shiga Toxin (e.g. STEC/VTEC) during human disease can produce a severe kidney pathology termed hemolytic uremic syndrome (HUS), and in severe cases systemic neuropathology (10, 11).

The coordinated regulation of T3SS expression and subsequent effector translocation are considered critical factors for EHEC/EPEC infection and disease progression (12–15). A genetic pathogenicity island, the locus of enterocyte effacement (LEE-PAI), encodes the structural protein components of the T3SS, along with regulators, chaperones and effectors (9). Additional pathogenicity islands found within the EHEC and EPEC chromosomes encode effectors that are secreted by the LEE encoded T3SS (16–19). Interestingly, a temporal hierarchy for translocon protein secretion (EspA, B and D) and effector secretion has been observed (20–22), indicating that EPEC has a tiered mechanism to first coordinate translocon and subsequently effector injection into host cells. Low calcium sensing by EPEC has been implicated in transitioning from translocon to effector protein secretion (20). The first secreted, or hierarchical effector, is the <u>t</u>ranslocated <u>i</u>ntimin <u>r</u>eceptor (Tir) (23, 24), although the mechanism behind this hierarchy remains elusive. Once Tir is translocated and localized within the host cytoplasmic membrane, it acts as a high affinity surface exposed receptor for intimin which is embedded within the outer membrane of EHEC/EPEC (25, 26). In concert with other host proteins, Tir further acts to subvert host cytoskeletal function. Notably, Tir has been shown to be essential for intestinal disease (5, 8), an observation consistent with the importance of Tir effector hierarchy. Several other effectors are involved in altering innate immune activity and signalling (EspF, NleA, NleB, NleC, NleD, NleE, and NleH1/H2) while others have subversive effects on cellular pathways (Map, EspH, NleG, reviewed in (3, 9)).

CesT is a 156 amino acid EPEC/EHEC multicargo <u>T</u>ype <u>3</u> <u>S</u>ecretion <u>C</u>haperone (T3SC, class 1B) that forms homodimers within the bacterial cytoplasm (27, 28). The CesT protein sequence (Genbank accession number CAS11488.1) is highly conserved (96-100% identity) across all serotypes and strains within publicly accessible databases. Located within the LEE-PAI, the *cesT* allele is transcriptionally controlled by two differentially regulated promoter elements (29). CesT homodimers have been shown to interact with at least 10 of the 21 annotated EPEC effectors (Tir, Map, EspF, EspH, EspG, EspZ, NleA, NleG, NleH1, and NleH2), which suggests a central role for this class 1B multicargo chaperone in the EPEC pathogenic strategy (27). CesT-effector interactions are believed to be mediated by a modestly conserved ‘chaperone binding domain’ (CBD) located within the amino terminal region of many effectors (23). Some EPEC/EHEC effectors apparently do not interact with CesT (e.g. NleC, NleD and NleE), and as expected many of these effectors have been experimentally demonstrated to not require CesT for secretion or translocation into host cells during bacterial infection (21).

CesT shares structural similarity with other T3SC of other pathogens, however it possesses a unique ~25 amino acid carboxyl terminal region. As a family of proteins, T3SC are thought to interact with the cytoplasmic face of the membrane located T3SS apparatus (22, 30). CesT has been shown to interact with EscN, a hexameric ATPase that is required for T3SS function (31, 32). A genetic screen to assess CesT function identified many CesT C-terminal substitution variants that were defective for effector secretion (24). Specifically, some CesT variants only supported Tir secretion and not other effectors, thus implicating the C-terminal region of CesT in coordinating effector secretion. Interestingly, in EHEC O157:H7, CesT has been shown to be tyrosine phosphorylated within its C-terminal region on residues Y152 and Y153 (33) and we have shown this for EPEC CesT as well (Runte and Thomas, unpublished results). Exactly how the multicargo CesT chaperone coordinates multiple effector secretion remains unknown.

In this study, we provide evidence for genetically separable functions linked to two strictly conserved C-terminal tandem tyrosine residues within a putative disordered region of CesT. We demonstrate a CesT mediated regulatory role during EPEC infection that impacts overall pathogenesis. The finding of functionally relevant and strictly conserved tandem tyrosines, each with differential roles in type 3 secretion associated events, is a novel paradigm in EPEC pathogenesis.

## Results

### CesT Tyr152 and Tyr153 are strictly conserved residues within a structurally disordered region

An EHEC CesT crystal structure has previously revealed a stable dimer (28). The C-terminal ~20 amino acids are located outside the dimerization interface, and the last 11 amino acids were not resolved within protein crystals, suggesting it might be a disordered terminal region. Moreover, this terminal region of CesT and specific amino acid residues within it, are important for effector secretion (24) (Fig 1A). CesT is highly conserved over its entire 156 amino acid sequence, displaying 96-100% identity for over 300 NCBI database entries (Dataset S1). This analysis included a diverse set of A/E pathogen serotypes from isolates obtained on multiple continents, with many associated with animal reservoirs (cow, pig, dog, rabbit, sheep). Low frequency conservative substitutions occur across most of the CesT sequence, however we noted amino acids Tyr152 and Tyr153 are without exception strictly conserved in all LEE-PAI pathogenic strains, including *Citrobacter rodentium*, an A /E mouse pathogen (Fig 1A). Notably, Tyr152 and Tyr153 of CesT have been observed to be phosphorylated in EHEC O157:H7 (33), and we have identified the same site-specific modifications for EPEC (Runte and Thomas, unpublished results) (Fig S6). These findings further suggest that this localized region is involved in secretory events. Monospecific antibodies raised against an EPEC CesT C-terminal peptide detected CesT proteins in EHEC O157:H7 and *C. rodentium* lysates, thus verifying native expression of the respective proteins (Fig 1B). The observation that two adjacent tyrosines are strictly maintained and are subject to post-translational modification within a structurally disordered region of CesT, strengthened our hypothesis that these specific tyrosine residues could be important for CesT function.

**Fig 1.**
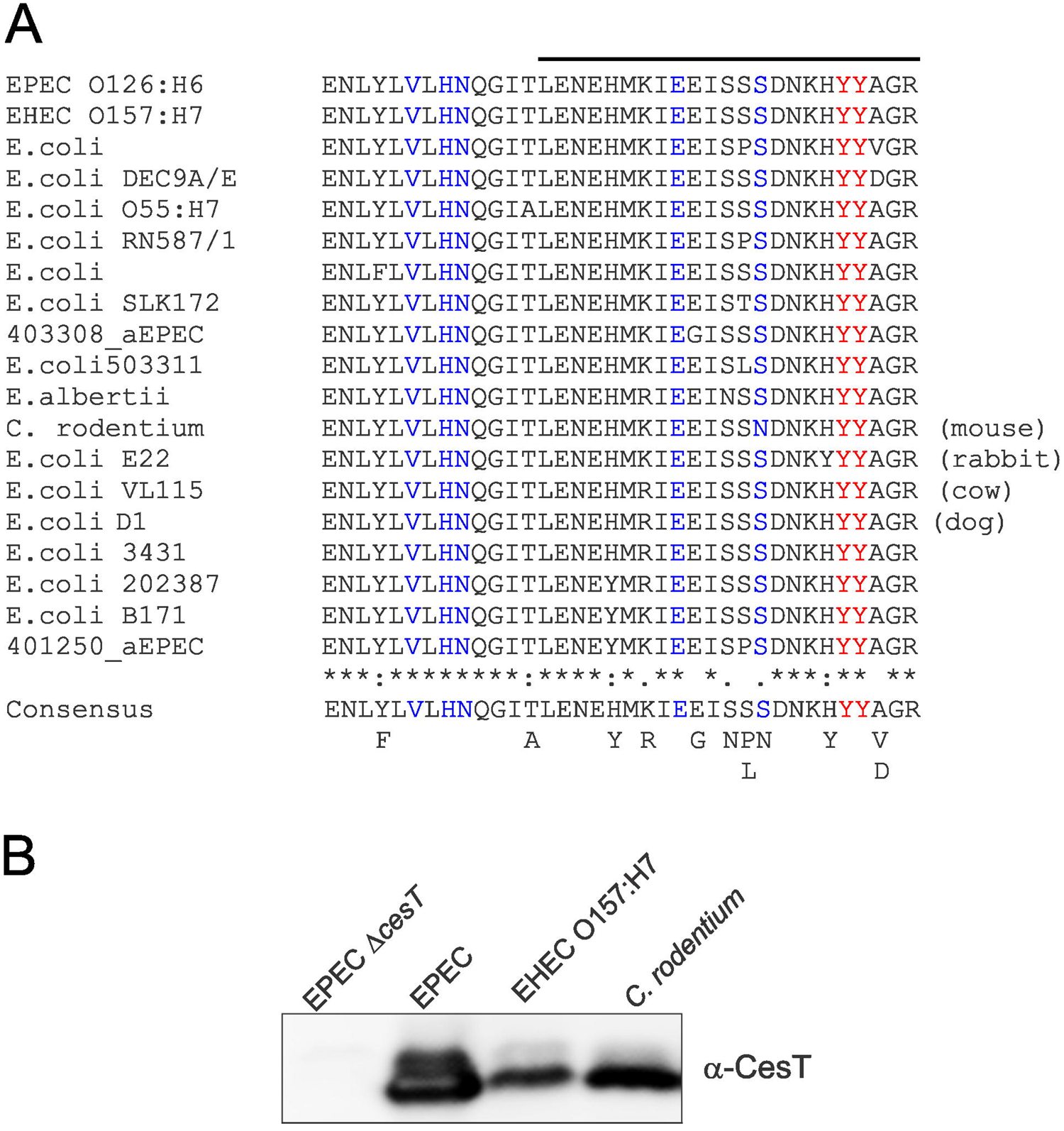
The C-terminal region of CesT is highly conserved in multiple clinical isolates, with strictly conserved tyrosines at positions 152 and 153. **(A)** Multiple sequence alignment of the C-terminal region of CesT. A consensus sequence is shown. Asterisks indicate invariant residues, and other observed amino acids at specific positions are identified. Amino acids labeled in blue have been implicated in effector secretion efficiency. The invariant tyrosine residues that are phosphorylated are shown in red. The black line above the EPEC sequence represents a peptide that was used to generate monospecific antibodies that were used for the blot in panel B. **(B)** Immunoblot of whole cell lysates from the indicated bacterial strains using anti-CesT antibodies directed against the C-terminal region of EPEC CesT.

### Efficient F-actin pedestal formation requires CesT Tyr152 or Tyr153

The Tir effector is an essential host colonization factor (5, 8). During infection of *in vitro* cultured human epithelial cells, CesT is required for efficient Tir translocation by EPEC (23). To investigate whether conserved CesT tyrosine residues contribute to Tir translocation, we performed HeLa cell infection assays with EPEC strains expressing CesT with conservative tyrosine to phenylalanine substitutions. This approach allows for retention of CesT secondary structure, and abrogates any potential for phosphorylation (i.e. removal of critical hydroxyl group). We generated EPEC strains that express i) CesT Y152F, ii) CesT Y153F, and iii) CesT Y152F, Y153F (non-phosphorylatable at positons 152 and 153). Importantly, these strains maintain the normal chromosomal operon organization that is involved in dual transcriptional regulation of *cesT* mRNA expression (29) and were found to produce the respective CesT variants (tyrosine to phenylalanine) at levels equivalent to wild type CesT as determined by immunoblotting with anti-CesT antibodies (Fig 2A). Therefore, CesT tyrosine residues 152 and 153 are not essential for maintaining CesT steady state levels in EPEC. Moreover, and as expected, these strains were unchanged for secreted levels of T3SS translocator proteins EspA, EspB, and EspD, along with EspC (type V secreted) (Fig 2A), a process known to be CesT-independent (31, 34).

**Fig 2.**
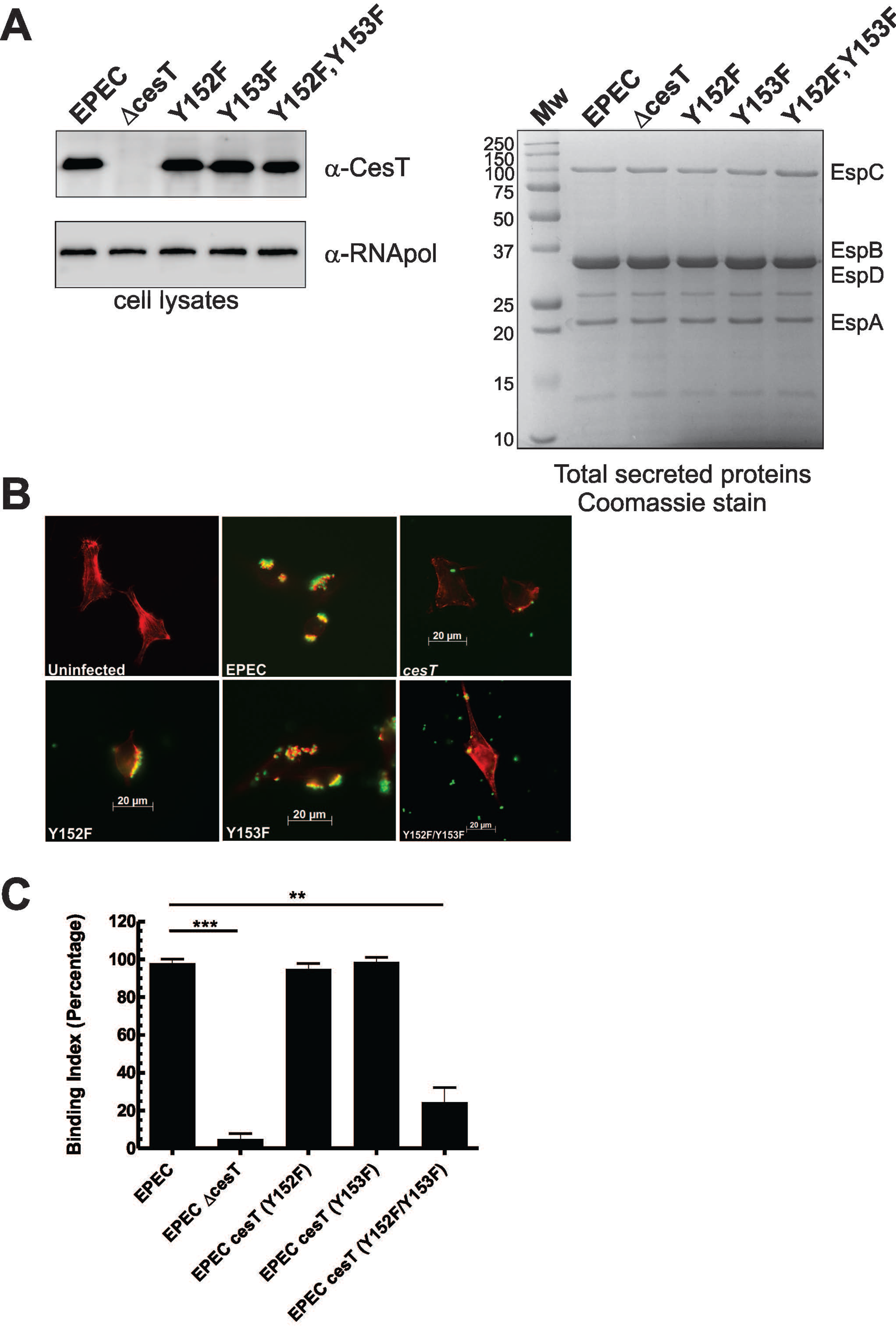
*In vitro* secretion and infection assays to characterize EPEC or EPEC mutants with tyrosine to phenylalanine substitutions within CesT. **(A)** Assessment of whole cell lysates and total secreted proteins derived from various EPEC strains. Whole cell lysates were probed with anti-CesT or anti-RNA polymerase antibodies. Total secreted proteins collected from the same cultures were stained with Coomassie G-250. The secreted protein profile of EPEC has been well characterized and the labeled proteins have been previously identified by mass spectrometry (23, 31). **(B)** Mock treated (uninfected) or EPEC infected Hela cells were stained with fluorescent phalloidin to detect F-actin (red). All bacterial strains contained a plasmid encoding green fluorescent protein. Note the appearance of robust F-actin condensation (actin pedestals) for specific bacterial strains. Representative images of each condition are shown. **(C)** Quantitation of EPEC strain intimate attachment using a defined binding index assay as described in the Methods section. Scale bar represent standard deviation from the mean. A paired t-test was used to determine statistical significance; ** p=0.0022, *** p=0.0001.

The ability of the CesT tyrosine to phenylalanine substitution variants to support A/E lesion formation was assessed using an *in vitro* HeLa cell infection assay where EPEC clusters HeLa cell F-actin in a Tir-dependent manner, forming actin-rich ‘pedestals’. We used fluorescence microscopy with a defined rule set to enumerate pedestals per HeLa cell for each infection condition (see methods). As expected, EPEC formed robust F-actin pedestals upon HeLa cell infection whereas in sharp contrast a *cesT* null mutant was significantly deficient in the process (Fig 2B). These data confirm previous reports and demonstrate that efficient EPEC actin pedestal formation is CesT-dependent (23, 34, 35). When EPEC expressing CesT Y152F or CesT Y153F was used to infect HeLa cells, F-actin pedestals were readily detected, resembling wild type EPEC. In contrast, when EPEC expressing non-phosphorylatable CesT Y152F, Y153F was used to infect HeLa cells, F-actin pedestal formation was significantly attenuated (p=0.002) (Fig 2C). We confirmed that each of the bacterial strains expressed their respective CesT variant by collecting infected HeLa lysates and immunoblotting for CesT (Fig S2). We also considered the possibility that CesT tyrosine substitution variants i) Y152F, ii) Y153F, and iii) Y152F,Y153F may not bind Tir efficiently. Using an established *in vitro* effector binding assay, it was determined that the CesT tyrosine to phenylalanine variants bound Tir (and other effectors) relatively equal to native CesT (Fig S3). Collectively, these data suggest that the CesT tyrosines (Y152 or Y153) contribute to the ability of EPEC to form F-actin pedestals on HeLa cells.

### EPEC expressing site specific CesT variants produce altered secreted levels of multiple effectors

Given our findings implicating conserved CesT C-terminal tyrosines for Tir injection into host cells, we set out to investigate if the secretion and translocation of other EPEC effectors are dependent on CesT tyrosine 152 and 153. Five CesT-interacting effectors (EspF, NleA, NleH1, NleH2, and Tir) were selected to generate chromosomally encoded C-terminal beta-lactamase (TEM-1) translational fusions within EPEC, Δ*cesT*, and each of the tyrosine to phenylalanine substitution mutants. The rationale for selecting these effectors was because they are encoded from three separate prophage or pathogenicity islands (LEE, PP2, PP6) and would thus widely inform on a potential role for the specific CesT tyrosine residues. This experimental approach allowed for detection of i) multiple effectors during *in vitro* secretion assays using high affinity commercial antibodies and, ii) quantitative real-time measurement of effector translocation during infection (i.e. effector injection into EPEC infected HeLa cells).

Initially, we assessed if the encoded effector-TEM-1 fusions were secreted into culture supernatants. All wild type EPEC strains supported efficient secretion of each effector-TEM-1 fusion (Fig 3A). The secretion of the effector-TEM-1 fusions were CesT-dependent in all but one case (EspF, see discussion), as they were not detected in culture supernatants derived from *cesT* null mutants (Fig 3B-F). For NleA-TEM-1, its secretion was abolished with substitution of CesT Y152 in two separate strains (Y152F, and Y152F, Y153F) (Fig 3A). Interestingly, in the case of *cesT* Y153F (where only CesT Y152 remains) NleA-TEM-1 secretion was significantly higher than wild type (Fig 3C; p=0.036). This represented a unique situation among the studied effectors so we assessed total cell lysates from the relevant strains which revealed that NleA-TEM-1 was exclusively detected when CesT Y152 was intact (only for wild type EPEC and *cesT* Y153F) (Fig 3G). Moreover, whole cell lysate levels of NleA-TEM-1 were consistently higher for *cesT* Y153F than WT EPEC, an observation that agreed with the secretion trends. Therefore, under these culture conditions, the data indicated that CesT Y152 was required for NleA expression, and that substitution of CesT Y153 to F153 increased the levels of NleA expression and its subsequent secretion.

**Fig 3.**
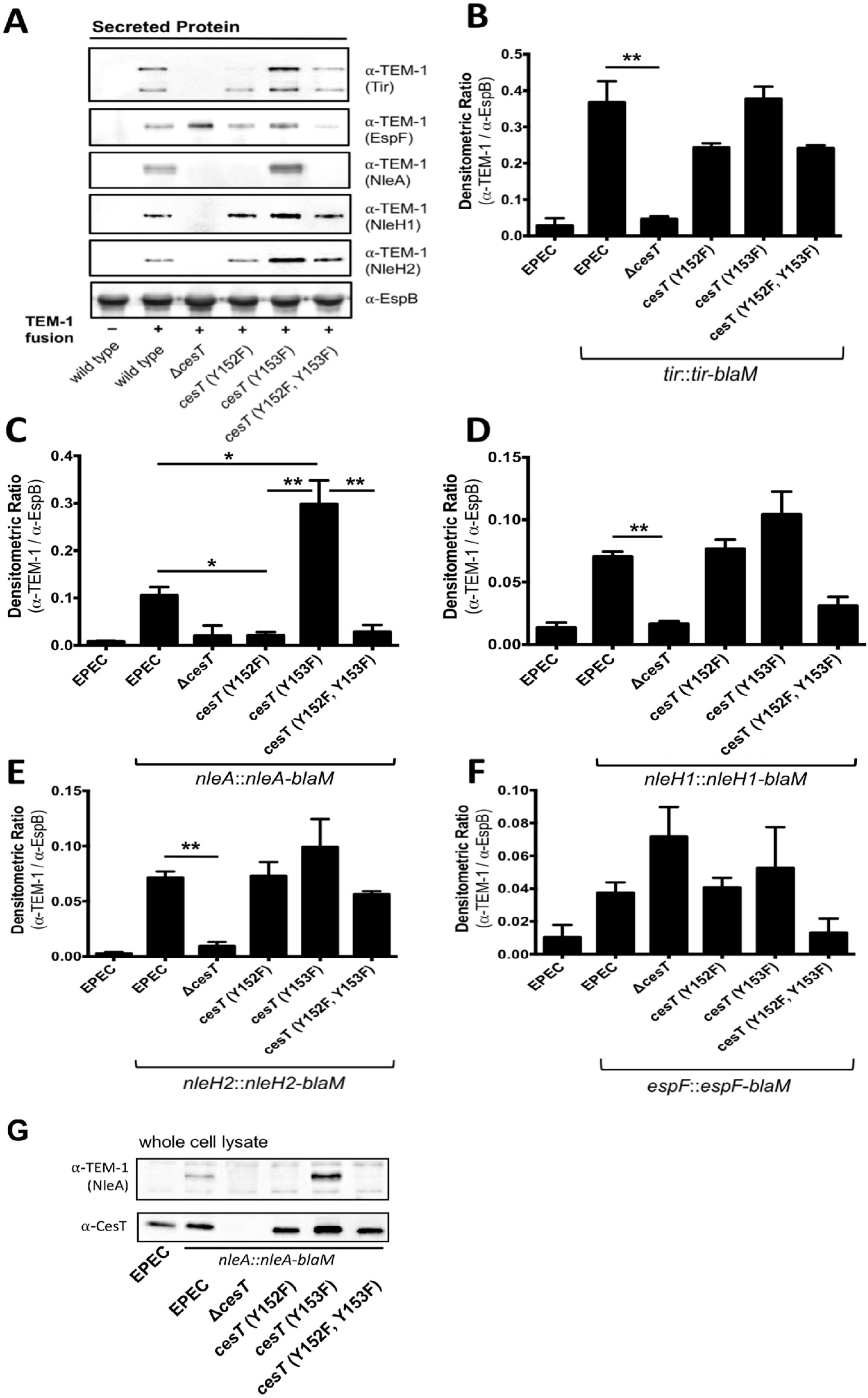
Characterization of effector-TEM-1 secretion in the context of chromosomal *cesT* variants. Each chromosomal *cesT* variant expresses a single monocopy effector-blaM fusion construct from its native promoter region. **(A)** The secreted protein fraction of each fusion construct and the parent EPEC strain, cultured in DMEM +2mM EGTA, normalized to OD_600_, was analyzed by immunoblotting. EspB (T3SS translocon protein) served as a loading control within each individual sample. Representative immunoblots from two independent experiments are shown. **(B-F)** Densitometry analysis of each immunoblot was normalized to EspB, and is quantitated as the ratio of effector-TEM-1 signal / EspB signal. Error bars represent standard deviation; data was compared by multiple unpaired t-test (* p < 0.05, ** p < 0.02). **(G)** Whole cell lysates of the indicated *nleA::nleA-blaM* EPEC strains were cultured in DMEM +2mM EGTA, normalized to OD_600_, and analyzed by immunoblotting.

Observed differences in Tir-TEM-1, NleH1-TEM-1 and NleH2-TEM-1 secretion in context of the CesT variants were less striking than for NleA-TEM-1. Nonetheless, secretion of Tir, NleH1, and NleH2 was always most efficient with Y152 intact (Fig 3A, 3D, and 3E). Furthermore, all EPEC strains expressing non-phosphorylatable CesT Y152F, Y153F demonstrated lower secreted levels of these effector-TEM-1 fusions than EPEC with native CesT. In each case total cell lysate levels of the Tir, NleH1, NleH2 effector-TEM fusions were comparable (data not shown). The results indicate that Y152 or Y153 are not absolutely required for *in vitro* secretion of Tir, NleH1, and NleH2, however, CesT Y152 promotes more efficient secretion of these effectors. The effector-TEM-1 secretion results are suggestive of an important role for CesT Y152 in overall EPEC effector secretion.

One disadvantage of studying effector-β-lactamase fusion proteins is that they represent altered and unnatural secretion substrates, which could potentially have different secretion outcomes. To address this, we employed an EPEC Δ*sepD* strain that is known to hypersecrete multiple natively expressed effector proteins into culture supernatants in a CesT-dependent manner (36) (Fig S4). We reasoned that this EPEC strain should exhibit site specific tyrosine dependent phenotypes associated with CesT (i.e. that agree with observations for strains expressing recombinant effector-TEM-1 fusions). We found that plasmid encoded CesT Y152F failed to restore total effector secretion, whereas CesT Y153F completely returned effector secretion. Therefore, data from two independent approaches indicated a critical role for Tyr152 of CesT in supporting total *in vitro* effector secretion for EPEC.

### CesT tyrosine to phenylalanine variants exhibit altered effector translocation kinetics during real-time *in vitro* infection conditions

The injection of EPEC effectors into host cells via the T3SS is a functionally relevant process leading to host cell subversion during infection. Therefore, we set out to investigate the role of the implicated CesT tyrosine residues in supporting the injection of NleA-TEM-1, NleH-TEM-1, NleH2-TEM-1, EspF-TEM-1, and Tir-TEM-1 into HeLa cells during EPEC infection. An empirically determined multiplicity of infection of 200 was used for these assays, which lead to notable translocation efficiency differences between EPEC and Δ*cesT* strains for all studied effector-TEM-1 fusions (Fig S5). For EPEC expressing Tir-TEM-1, translocation was detected 20-30 minutes post infection reaching its maximal level at 60-70 minutes post-infection (Fig 4A). Here, Tir-TEM-1 translocation was found to be significantly different from a *cesT* null mutant that supported near baseline levels of Tir-TEM-1 translocation (p<0.01) (Table S1). Analyses of specific EPEC CesT tyrosine to phenylalanine substitution variants revealed efficient Tir-TEM-1 translocation for bacteria expressing CesT Y152F or CesT Y153F, that closely resembled EPEC with normal CesT. In contrast, EPEC expressing non-phosphorylatable CesT Y152F, Y153F consistently supported reduced levels and a slower rate of Tir-TEM-1 translocation (see curve slope in Fig 4A). The reduced levels did not reach statistical significance compared to EPEC expressing CesT, however indicate a trend implicating CesT residues Y152 and Y153 in Tir translocation efficiency (see F-actin pedestal assay observations (Fig 2). In assessing this data, it should be noted that enzymatic translocation and F-actin pedestal assays measure different biological processes. Specifically, Tir-TEM-1 injection into host cells is measured by the enzymatic activity of TEM-1 on a cell-loaded CCF2-AM substrate (21, 37), whereas F-actin pedestals are the outcome of a multi-step host driven signalling cascade initiated by efficient Tir injection and clustering.

**Fig 4.**
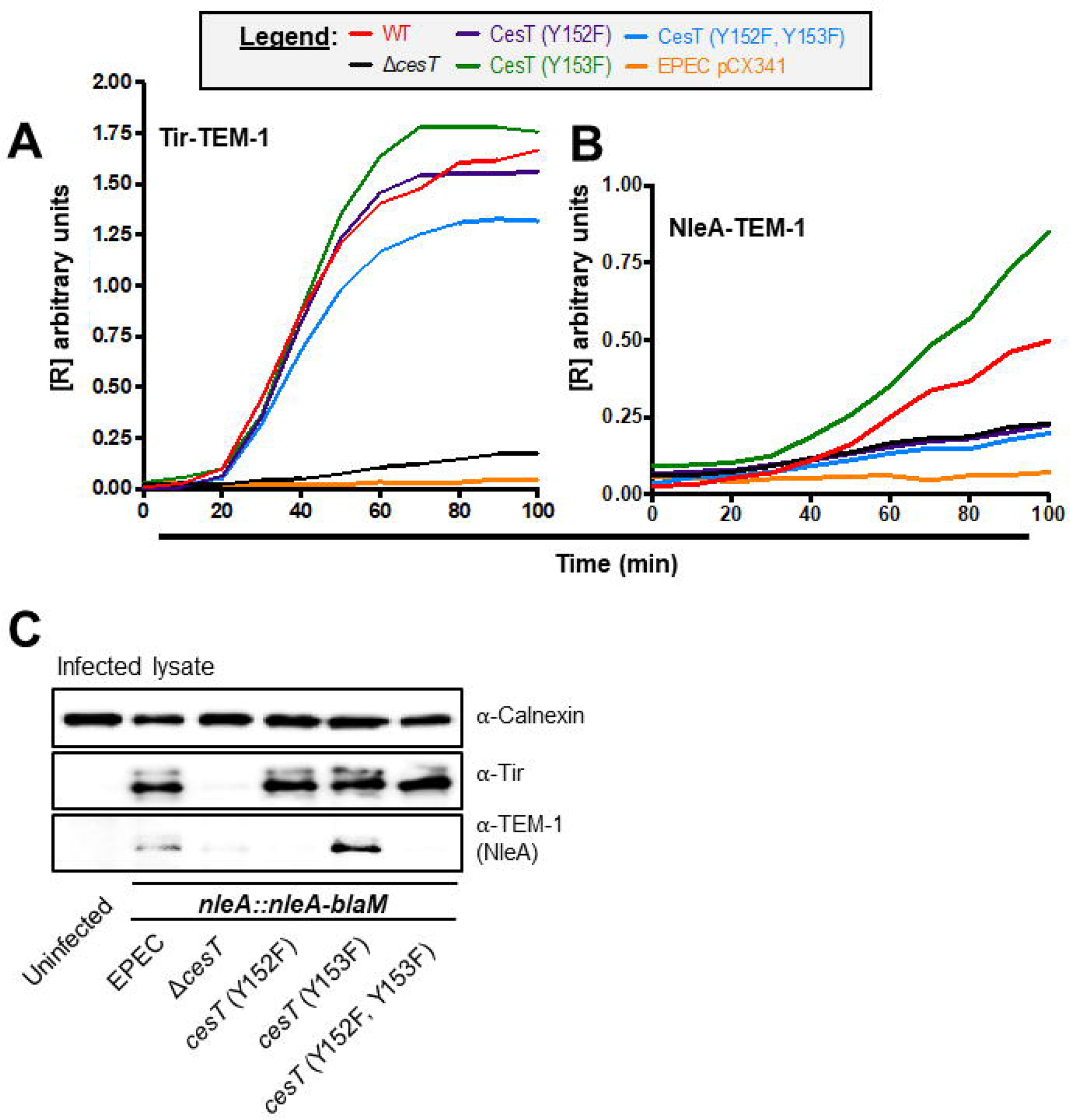
Tyrosine to phenylalanine amino acid substitutions within the CesT C-terminal domain impact NleA translocation and expression. EPEC and *cesT* variant EPEC strains, each containing a chromosomally encoded effector-*blaM* gene, were subjected to kinetic translocation analysis for Tir **(A)** or NleA **(B)**. HeLa cells were preloaded with CCF2/AM and infected at MOI 200 with bacterial strains ‘pre-activated’ in DMEM for 4 hours. The [R] value (y-axis) is a quantitative ratio representative of effector-TEM-1 accumulation within infected HeLa cells. Wild type EPEC carrying pCX341 (expresses native, unfused TEM-1) was used as a negative control for non-T3SS-specific beta-lactamase translocation. The data is representative of three independent infection experiments carried out in duplicates. For data presentation clarity, error bars are omitted and statistically significant differences are noted in S2 Table 2. **(C)** Total lysates were collected after 2h from mock infected or HeLa cells inoculated at MOI 200 with the indicated *nleA::nleA-blaM* strains pre-activated in DMEM. Calnexin served as a loading control for immunoblot analysis.

Next, we assessed the role of CesT tyrosines on NleA-TEM-1 translocation into Hela cells. EPEC expressing CesT exhibited augmented NleA-TEM-1 translocation 50 minutes post infection deviating from the baseline levels observed for the *cesT* null strain (Fig 4B). Interestingly, EPEC expressing CesT Y153F resulted in an increased amount of translocated NleA-TEM-1 at 40 minutes post infection, outperforming EPEC with native CesT. This reached statistical significance by 80 minutes post infection (Table S2). In sharp contrast, EPEC expressing CesT Y152F or CesT Y152F, Y153F produced NleA-TEM-1 translocation levels that mimicked baseline Δ*cesT* levels. We verified that these strains were translocation competent (capable of injecting effectors) by assessing Tir translocation into host cells by immunoblotting the respective infection monolayer lysates. Indeed, EPEC expressing CesT Y152F or CesT Y152F, Y153F were shown to inject Tir into host cells (Fig 4C), indicating that a functional T3SS was expressed. Collectively, these data suggest that CesT Y152 was required for efficient NleA translocation and that increased translocation levels were achieved when CesT Y153 was substituted to phenylalanine.

EPEC strains expressing CesT Y152F, or CesT Y152F, Y153F were observed to support reduced translocation levels for NleH1-TEM-1, and NleH2-TEM-1 relative to EPEC with native CesT (Fig 5A and 5B). While translocation efficiency for these effectors was not reduced to Δ*cesT* levels (as noted for NleA-TEM-1), statistically significant differences were reached at 70-100 minutes post infection in comparison to EPEC expressing CesT (summarized in detail, Table S2). For EPEC expressing CesT Y153F, NleH1-TEM-1 and NleH2-TEM-1 late translocation levels (80-100 minutes postinfection) were slightly elevated compared to EPEC expressing CesT although the difference did not reach statistical significance. In the strains expressing EspF-TEM-1, EPEC with CesT Y153F supported the highest translocation level for all strains evaluated, even surpassing EPEC with native CesT (Fig 5C, EspF represents an unusual effector in that it interacts with two different chaperones, CesF and CesT, see discussion). In summary, these data suggest that EPEC expressing CesT with Y152 intact supported high translocation efficiency for NleH1 and NleH2, although Y152 was not strictly required for translocation of these effectors. EspF translocation appeared to be modestly enhanced when CesT Y153 was substituted to phenylalanine.

**Fig 5.**
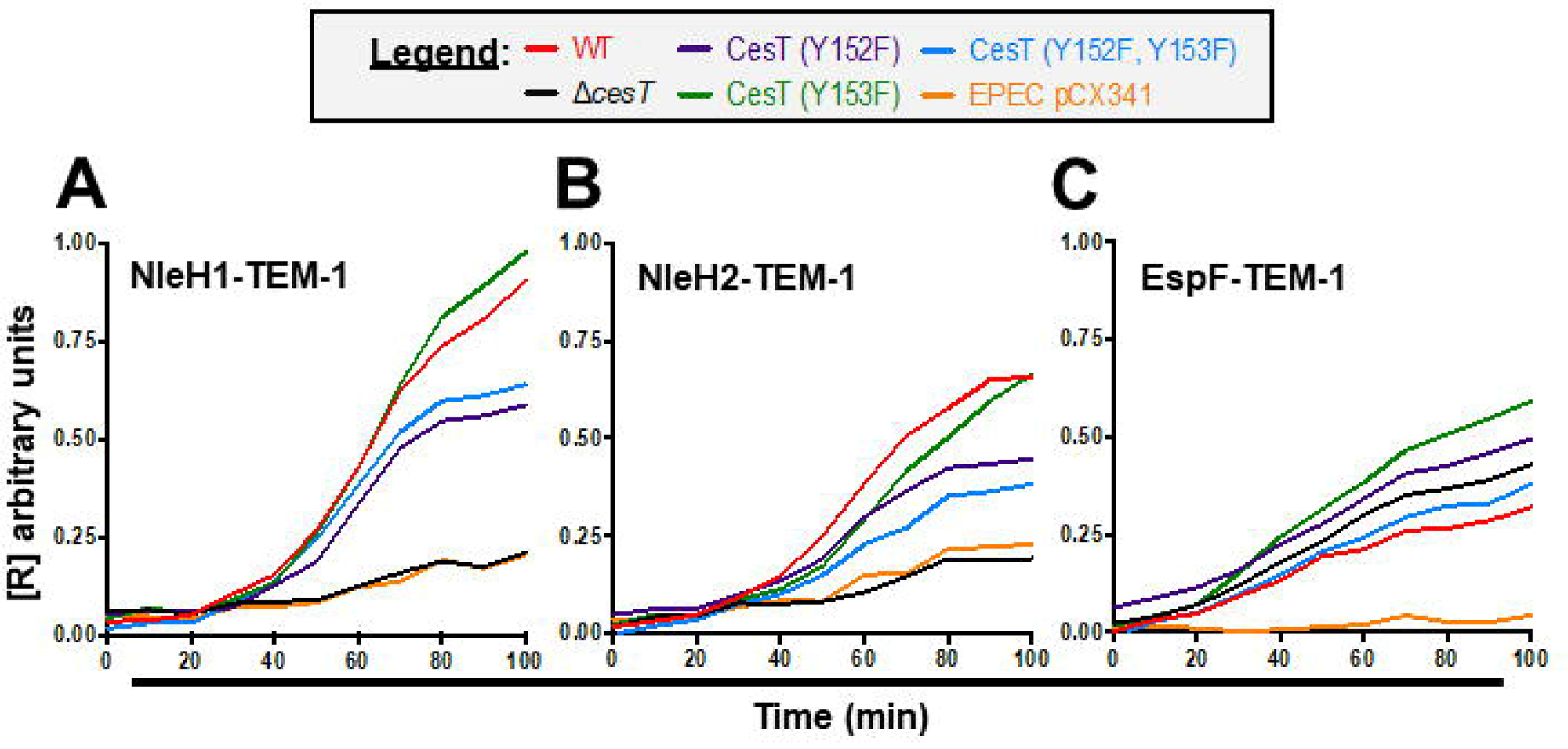
Site-specific amino acid substitutions within the CesT C-terminal domain impact effector translocation efficiencies. EPEC and *cesT* variant EPEC strains, each containing a chromosomally encoded effector-*blaM* gene, were subjected to kinetic translocation analysis for NleH1 **(A)**, NleH2 **(B)**, and EspF **(C)**, The [R] value (y-axis) is a quantitative ratio representative of effector-TEM-1 accumulation within infected HeLa cells. Wild type EPEC carrying pCX341 (expresses native, unfused TEM-1) was used as a negative control. The data is representative of three independent infection experiments carried out in duplicates. For data presentation clarity, error bars are omitted and statistically significant differences are noted in Table S2.

### *C. rodenrtium* requires the presence of CesT Tyr152 and Tyr153 for efficient mouse colonization

We set out to investigate whether specific C-terminal CesT tyrosines are important for bacterial intestinal colonization and disease in a natural mouse infection model for A/E pathogenesis. *C. rodentium* has been widely used by researchers to assess the role of its putative virulence factors in mice, including many aspects of its cognate T3SS and numerous effectors (8, 38–41). Importantly *C. rodentium cesT* null mutants have been shown to be strongly attenuated for mouse intestinal colonization and disease (8).

Given the important role for the CesT Y152 for EPEC effector injection into host cells, we generated *C. rodentium* substitution mutants that express either CesT Y152F, CesT Y153F, or CesT Y152F, Y153F. Next, 6 groups of healthy C57BL6 mice (n=5) were infected by oral gavage with the following *C. rodentium* strains; i) wild type, ii)*cesT* Y152F, iii) *cesT*Y153F, iv) *cesT*Y152F, Y153F, v)Δ*cesT*, and vi)Δ*nleA*. The last 2 groups were included to provide a comparison to previously published observations for highly attenuated *C. rodentium* mouse infections (8, 42). A seventh group of mice was kept uninfected and monitored throughout the course of the 9 day experiment. Fecal shedding of *C. rodentium* was followed for each group of mice at day 3, 5, 7, and 9. Mice were then sacrificed at Day 9 to assess *C. rodentium* colon counts. Importantly, no *C. rodentium* were detected at any time in the uninfected mice (data not shown).

Three days post-infection, most mice from all treatment groups except Δ*cesT* shed *C. rodentium* at levels clustering at 10^5^ cfu/g feces although considerable variability in shedding was observed for individual mice in each group (Fig 6A). For 4/5 Δ*cesT* infected mice, *C. rodentium* was not detected in fecal samples, with the remaining mouse shedding ~10^5^ cfu/g feces. Notably, one and two mice from wild type and *cesT* Y153F respectively shed at ~10^7^ cfu/g feces. At day 5, average *C. rodentium* counts in feces were markedly different among groups with wild type and *cesT* Y153F having the highest average values near ~10^8^ cfu/g feces (Fig 6A). The average count for *cesT* Y152F was lower at ~10^7^ cfu/g feces, followed by an even lower average count of ~10^5^ cfu/g stool for *cesT* Y152F, Y153F. At or near Day 7, C57BL6 mice are known to exhibit maximal *C. rodentium* fecal shedding (43). As expected, the average *C. rodentium* fecal shedding count for wild type was high at ~10^8^ cfu/g stool (Fig 6A). Similar average values for *cesT* Y152F, *cesT* Y153F, and *cesT* Y152F, Y153F were also detected. In contrast, and as expected, Δ*nleA* and Δ*cesT* strains were detected at much lower averages of ~3.0×10^6^ and 3.0×10^4^ cfu/g feces respectively. At day 9, fecal *C. rodentium* average counts for wild type, *cesT* Y152F, and *cesT* Y153F remained high at ~10^8^ cfu/g stool (Fig 6A). Critically, the fecal shedding average count for *cesT* Y152F, Y153F sharply declined to 2.8×10^6^, close to the average detected for Δ*nleA*. Lastly, Δ*cesT* exhibited the lowest average fecal count at 4.8×10^4^.

**Fig 6.**
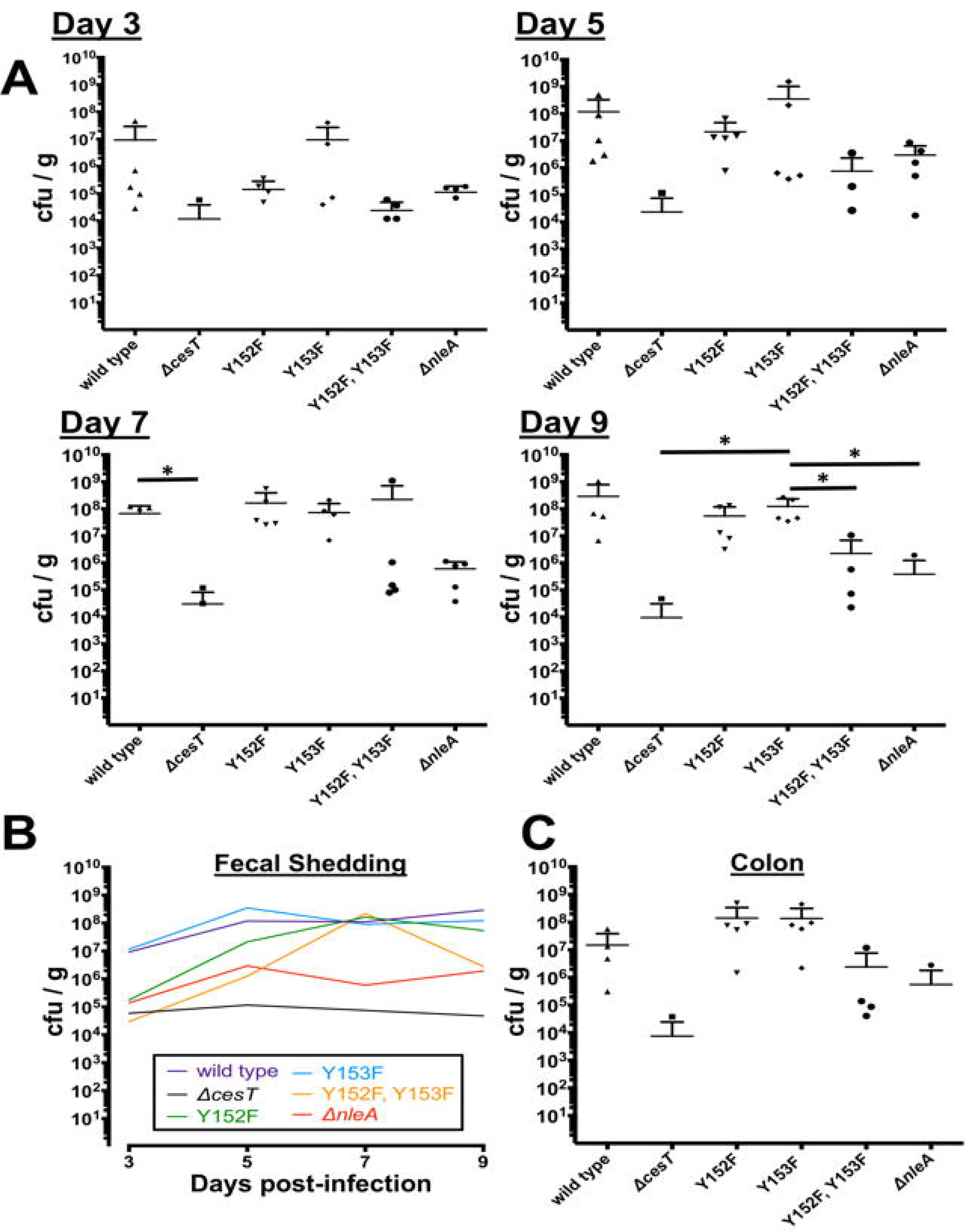
Shedding of *C. rodentium* strains determined from fecal samples of infected C57BL/6 mice (n=5 per group). Bacterial enumeration from feces collected at **(A)** day 3, day 5, day 7, and day 9 are represented with lines to indicate the mean+ + SD. **(B)** The dynamics of shedding is presented as the mean for each experimental group at each time point. **(C)** Bacterial enumeration from excised colon tissue at day 9 postinfection. Statistically significant cfu/g differences were determined by multiple unpaired t-tests where indicated by asterisks (*, P < 0.05).

Taken together, the data indicate an early intestinal colonization defect for *C. rodentium cesT* Y152F and *cesT* Y152F, Y153F relative to wild type *C. rodentium*, a trend that was evident up to day 5 post infection. As the infection progressed, *cesT* Y152F, Y153F were shed at high numbers but rapidly declined, differing from wild type, *cesT* Y152F, and *cesT* Y153F bacteria which maintained high fecal shedding counts. The fecal shedding data (summarized in Fig 6B over 9 days) suggest that CesT Y152 and Y153 jointly contributed to efficient colonization of the mouse intestine although the presence of a single tyrosine at Y153 (i.e. strain *cesT* Y152F) was sufficient to support delayed colonization.

Next, on Day 9 the mice were sacrificed and colon samples were collected to assess tissue and mucosal-associated *C. rodentium* counts from each group of mice. As shown in Figure 6C, mice infected with wild type, *cesT* Y152F, and *cesT* Y153F each had high average colon counts, ranging from 1.5×10^7^ (wild type) to 1.4×10^8^ cfu/g tissue. Consistent with reduced fecal counts, mice infected with *cesT* Y152F, Y153F exhibited a lower average colon burden at 2.4×10^6^. As expected, mice infected with Δ*nleA* or Δ*cesT*were highly attenuated and had a very low colon tissue burden ranging from 10^4^ to 10^6^ with only 1 mouse from each group producing counts at day 9.

## Discussion

In this study, we provide evidence suggesting that strictly conserved tandem tyrosines within CesT function to regulate type 3 effector secretion within A/E pathogens. Using complementary genetic, biochemical, and infection biology approaches, we implicate CesT Y152 and Y153 in the development of enteric disease. Strikingly, bacteria expressing structurally conservative phenylalanine substitutions in place of specific tyrosines within CesT, were significantly deficient in effector translocation and strongly attenuated for *in vivo* colonization. Another new observation for the T3SS chaperone CesT is the dependence of a specific tyrosine residue in the expression of an effector protein, in this case demonstrated for EPEC NleA.

CesT Y152 appears to be correlated with maximal effector translocation efficiency, a trend observed for multiple effectors found in different EPEC prophage and pathogenicity islands. The only exception we observed was for EspF translocation. EspF interacts with its own chaperone CesF (44), yet can also interact with CesT (23, 31). Curiously, we observed maximal levels of EspF translocation in context of CesT Y153F (normal for Y152). EspF thus represents an example of an effector with a very complex T3SS trafficking paradigm.

Our data implicates CesT Y152 as a critical node for efficient EPEC effector translocation. Evidence for this interpretation comes from multiple experiments. Firstly, EPEC CesT Y152F was shown to support significantly reduced levels of effector translocation for NleA, NleH1, and NleH2. Secondly, CesT Y152 was strictly required for NleA expression, which is a unique and novel observation among the effectors we evaluated. Thirdly, phenylalanine substitution at the adjacent CesT 153 residue generated enhanced translocation levels of many effectors, exceeding levels supported by wild type CesT. Critically, these observations raise the paradoxical question as to why A/E pathogens have not undergone selective evolutionary pressure to alter Tyr153 to Phe153 via mutation (a single nucleotide change). The lack of such an occurrence, given its apparent effector translocation benefits, strongly suggests that A/E pathogens maintain a complex regulatory interplay at adjacent tyrosines (Y152 and Y153), a feature that we find to be important for effector secretion.

It has been proposed that CesAB (EspA chaperone) and CesT interact with an inner membrane T3SS assembly in an EscV-dependent manner (22). Furthermore, a C-terminal tail of CesAB was critical for the EscV interaction. It is tempting to speculate the same is true for CesT, especially in context of our previous observations relating to C-terminal tyrosine residues in CesT (24). Interestingly, FlgN, a Salmonella type III export flagellar chaperone, requires a highly conserved C-terminal Tyr122 residue to interact with FlhA (its EscV homologue) to support flagellar assembly (45). Moreover, another multicargo type III secretion chaperone, HpaB of *Xanthomonas*, was shown to interact with HrcV (EscV family) (46). Therefore, it appears that chaperone ‘docking’ interactions with FlhA/EscV homologues is a recurring theme in type III secretion, and perhaps site specific and contextual tyrosines are involved.

CesT is known to be required for efficient *C. rodentium* mouse intestinal colonization (8). We have reproduced those results and now extend the findings by revealing C-terminal tandem tyrosine residues within CesT are important for early intestinal colonization. We did observe that *C. rodentium* (CesT F152, F153) did replicate within mice, but was cleared from the mouse colon more rapidly than wild type *C. rodentium*. Recently, it has been demonstrated that the establishment phase of *C. rodentium* pathogenesis *in vivo* is restricted to a very short window of opportunity (~18 hours) that determines intestinal intimate attachment and disease severity (41). Indeed, our data directly agree with those findings in that bacteria expressing CesT F152, F153 were attenuated for F-actin pedestal formation (EPEC) and mouse intestinal infection (*C. rodentium*). Therefore, CesT acts as a critical interaction node within the bacterial cell that supports A/E pathogenesis.

It has not escaped our attention that tyrosine can be phosphorylated in biological systems, the result which often creates high affinity bindings sites for other proteins. Indeed, a phosphotyrosine proteomic study with EHEC O157:H7 identified CesT as being monophosphorylated on Y152 or Y153 during *in vitro* growth conditions (33). We have also identified that EPEC phosphorylates CesT on Y152 or Y153 (Fig S6), thus demonstrating that this site-specific post-translational modification is present in another closely related A/E pathogen. Substitution of CesT Y152 to F152 would abrogate localized phosphorylation, hence the absence of site-specific phosphorylation might explain our observations. The same would be true for CesT F153. In the case of CesT F152, F153, absolutely no localized phosphorylation could occur. Strikingly, our experiments reveal that bacteria expressing non-phosphorylated CesT F152, F153 exhibited the lowest effector translocation efficiencies (*in vitro*) and delayed mouse intestinal colonization (*in vivo*). Alternatively, the hydroxyl group on tyrosine 152 and 153 (and absent from the structurally similar phenylalanine) might serve a very localized role within CesT such as hydrogen bonding with another protein. Critically, the current data cannot distinguish the aforementioned possibilities. The discovery of a kinase responsible for phosphorylating CesT along with numerous other experiments will be required to address these very challenging questions.

The prophage encoded effector NleA is a critical virulence determinant of A/E pathogens (47, 48)(Fig 6). EPEC NleA expression is regulated by a post-transcriptional process where *nleA* mRNA is bound by the RNA binding protein CsrA to prevent its translation until T3SS-mediated host cell contact (49). CesT was further shown to relieve translation inhibition by interacting and displacing CsrA from the *nleA* mRNA transcript. Here, we provide evidence that only CesT with Y152 intact supports NleA expression. Surprisingly, we observed that CesT Y153F (with Y152 intact) significantly outperformed wild type CesT for NleA protein expression levels and translocation kinetics. This raises the intriguing possibility that Y152 of CesT might serve as a critical site for its interaction with CsrA, and thus promote NleA translation.

T3SS class 1 chaperones have been historically considered a group of structurally conserved proteins that function to maintain effector proteins in a secretion competent state within the bacterial cell (50–52). Evidence now indicates that T3SS chaperones also function to recruit effectors either to a‘sorting platform’ or the cytoplasmic face of the inner membrane associated T3SS apparatus resulting in effector secretion (22, 53). For the multicargo chaperone CesT, effector binding and recruiting functions have been demonstrated to be functionally independent (24, 31), and here we implicate strictly conserved CesT tyrosine residues in differential regulation of effector secretion. We hypothesize that CesT Y152, and perhaps its phosphorylation, serves as a high affinity binding site for a specific protein component of the T3SS.

## Materials and Methods

### Bacterial Strains and Growth Media

Bacterial strains generated and used in this study are listed in Table S3. Bacteria were routinely cultured in Luria broth (LB) (1% [w/v] tryptone, 0.5% [w/v] yeast extract, 1% [w/v] NaCl) at 37°C. Antibiotics (Sigma) were added when appropriate, to a final concentration of 50 μg/ml kanamycin, 50 μg/ml streptomycin, 30 μg/ml chloramphenicol, 100 μg/ml ampicillin. For experiments that required activation of T3SS gene expression, bacteria were cultured without antibiotics, under static conditions in serum-free DMEM (Invitrogen Catalogue #11995) in a humidified, 5% CO_2_ incubator at 37°C.

### Isolation of Genomic and Plasmid DNA

Genomic DNA was isolated from bacterial strains using the Purogene genomic DNA isolation kit (Gentra systems). Plasmids were isolated from bacterial strains using the QIAprep spin miniprep kit (Qiagen).

### Construction of *cesT* substitution mutants in EPEC E2348/69 and *Citrobacter rodentium*

EPEC and *C. rodentium* strains that express CesT tyrosine to phenylalanine variants were generated by mutating the chromosomal *cesT* allele using allelic exchange (24). Briefly, primers NT379 and NT380 were used in a PCR with EPEC genomic DNA to generate a 857 bp DNA product that was directionally cloned into pBluescript SacI and EcoRV sites, to generate pEPcesT. This plasmid was digested with NdeI and KpnI to serve as the vector backbone for synthesized *cesT* sequence gene blocks (Integrated DNA Technologies) that code for specific tyrosine to phenylalanine substitutions within CesT (Table S4). The gene blocks were directionally cloned as NdeI/KpnI fragments. The respective recombinant clones which encompass the *tir-cesT-eae* locus (each containing specific *cesT* mutations) were then subcloned as SacI/KpnI DNA fragments into suicide plasmid pRE112 using DH5αλ*pir* as a cloning host. A similar approach was used for *Citrobacter* using primers NT381 and NT382 for PCR with *Citrobacter* DNA to generate pCRcesT, followed by ligation cloning to gene blocks coding for specific Citrobacter *cesT* alleles in pRE112. All *cesT* mutations within their respective suicide plasmid constructs were verified by DNA sequencing. Suicide plasmid constructs were mobilized into recipient strains via triparental mating with recipient single crossover merodiploid clones (EPEC or *Citrobacter*) being selected with solid growth media containing streptomycin and chloramphenicol (for EPEC) or ampicillin and chloramphenicol (for *Citrobacter*). After streak purification of merodiploid clones on selective antibiotic media, sucrose selection was performed to isolate mutants as previously described (24). Bacterial isolates that exhibited growth on sucrose but not chloramphenicol were then streak purified. For each targeted mutation, several isolates were subjected to PCR analysis for the *cesT* allele, followed by DNA sequencing of the PCR amplicons. This approach yielded several *cesT* mutants.

### Protein electrophoresis and Immunoblotting

All protein samples were separated by SDS-PAGE as previously described (54). For routine protein gels, separated polypeptides were visualized by Coomassie G-250 staining. Alternatively, for mass spectrometry analyses, colloidal blue or silver staining was employed. Pre-stained All-blue Protein Markers (Bio-Rad)) were routinely used as molecular weight standards. For immunoblotting, separated polypeptides were transferred to Immobilon-P membrane (Millipore), blocked with skim milk (5% [w/v] in Tris buffered saline+ Tween20 [TBS-T]), and then incubated with specific antibodies at working concentrations; anti-FLAG, 1:10000 (Sigma); anti-RNApol, 1:2000 (Santa Cruz Biotech); anti-calreticulin,1:2000 (Calbiochem); anti-TEM-1, 1:5000 (Quantum Biosciences); anti-Tir, 1:1000 (23); goat anti-mouse IgG conjugated to horse radish peroxidise (HRP), 1:5000 (Rockland immunochemicals); goat anti-rabbit IgG conjugated to HRP, 1:5000 (Rockland immunochemicals). High affinity anti-CesT monospecific antibodies were raised against a synthetic peptide (LENEHMKIEEISSSDNK) corresponding to the C-terminal region of CesT and were affinity purified against the peptide by the supplier (Pacific Immunology) and used in immunoblots at a 1:10000 dilution (24). Immunoblots were developed using Clarity chemiluminescence reagent (Bio-Rad) and data captured on a VersaDoc 5000MP (Bio-Rad). Densitometry analyses of chemiluminescent signals was performed using ImageLab software (BioRad). All immunoblotting experiments were performed three independent times, with a representative image shown.

### Hela cell infection and F-actin pedestal assays

HeLa cells (American Type Culture Collection) were seeded (5×10^5^) into 60 mm culture dishes and grown overnight in DMEM supplemented with 10% fetal bovine serum (Invitrogen). Monolayers were then infected with EPEC or EPEC mutant strains for 3.5 hours. The infected Hela cell monolayers were then mechanically fractionated as previously described (55). The resulting samples were prepared for SDS-PAGE and immunoblotting. For F-actin pedestal assays, Hela cells on glass coverslips were infected with EPEC strains for 3.5 hours (or 9 hours with *Citrobacter* strains). The samples were washed three times with PBS to remove weakly or non-adherent bacteria and then fixed with paraformaldehyde. The samples were then prepared for fluorescence microscopy as previously described (23). Quantification of intimate bacterial attachment (binding index) was conducted as previously described (56). Briefly, the percentage of HeLa cells harboring at least five GFP-positive bacteria (identified by GFP fluorescence) that were associated with F-actin condensation were quantified. Three independent experiments were performed, with at least 50 Hela cells examined per sample.

### Generation of Chromosomally Encoded Effector-β-lactamase Fusions

The coding regions of CesT-interacting T3SS effectors NleA, NleH1, and NleH2 were amplified by PCR and cloned into pCX341 to generate C-terminal translational fusions to TEM-1 beta lactamase (encoded by the *blaM* gene). This cloning step was carried out in DH5αλpir *E. coli*. The effector-*blaM* fusion constructs in pCX341-based plasmids were excised with KpnI and XbaI digests, and subcloned into the pCX391 suicide plasmid digested with KpnI and XbaI. These constructs were added to competent DH5αλpir cells and transformants selected on LB tetracycline (10μg/mL) plates. The pCX442 (Tir-TEM-1) and pCX446 (EspF-TEM-1) suicide plasmids derived from pCX391 were previously received from Ilan Rosenshine. These plasmids were transformed into DH5aA,pir to finalize a total of five donor strains to be used in a conjugation experiment. Each donor contained a single pCX391-based effector-*blaM* fusion that was to be targeted for chromosomal integration within five different EPEC recipient strains: *wild type*, *ΔcesT*, *cesT*(*Y152F*), *cesT*(*Y153F*), and *cesT*(*Y152F*,*Y153F*). Single crossover integration of the plasmid via homologous recombination conferred resistance to tetracycline for the EPEC recipients. Streptomycin (50 μg/ml) was included in the selection media to prevent growth of DH5α*λ*,pir donors. Suicide plasmid-encoded effector-*blaM* constructs were verified by sequencing prior to conjugation.

### Real-Time Effector Translocation Assays

On day 1, HeLa cells were seeded in 96-well plates (black with clear bottom, ThermoFisher Scientific) at a density of 2×10^4^ cells/well in DMEM (Invitrogen) supplemented with 10% fetal bovine serum (FBS). In parallel, bacterial strains encoding chromosomal *effector-blaM* fusions were grown overnight in LB at 37°C, shaking at 200 rpm. On day 2, bacterial cultures were diluted 1/200 into DMEM and grown statically in conditions known to stimulate T3SS expression (37°C, 5% CO_2_ for 4 hours to OD_600 nm_ of 0.2-0.35) to create a ‘pre-activated’ culture. At the 3-hour mark of the culture preactivation, HeLa cells were washed twice with <u>D</u>MEM +10% <u>F</u>BS +2.5 mM <u>p</u>robenecid (herein referred to as DFP media), and treated with 20 μL of 6X CCF2-AM substrate loading solution dissolved in 100μL DFP media. The cells were incubated for 40 min at room temperature in the dark, then washed an additional 3X with 100 μL DFP to titer out extracellular CCF2-AM substrate from the growth media. OD_600 nm_ measurements were recorded for each bacterial culture at the 4 hr mark of pre-activation to normalize infection inoculum to MOI 50, 100, or 200. Immediately upon bacterial inoculation, the plates were placed in a plate reader (Victor X5; PerkinElmer) set to 37°C, which was where the infection took place. Infected cells were excited at 405 nm, and emission was recorded at 460 nm and 535 nm at 10 minute intervals over the course of infection. Experimental data was collected with PerkinElmer 2030 software and adjusted for background blue and green fluorescence using the following formula in Microsoft excel (© 2015):

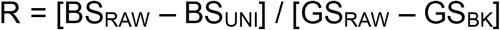

BS_RAW_, measured product fluorescence at 460 nm; BS_UNI_, average background 460 nm fluorescence from uninfected HeLa cells loaded with CCF2-AM substrate at each measured time-point; GS_RAW_, measured product fluorescence at 535 nm; GS_BK_, average background 535 nm fluorescence from uninfected HeLa cells without CCF2-AM substrate. To normalize well-to-well variation in number of HeLa cells and/or efficiency of CCF2-AM substrate uptake, all samples were run in duplicates, and R values for each condition at each time-point were averaged. Real-time translocation data is representative of average R values for at least three independent experiments. Wild type EPEC expressing β-lactamase (unfused) was included in each assay as an experimental negative control.

### Mouse infections with *Citrobacter rodentium*

Female C57BL/6 mice (Charles River Laboratories) were acclimatized for 7 days in our specific pathogen-free facility. The mice were then randomized into seven groups (n=5 per group), for six treatments and 1 mock infection. Bacterial inocula for the respective mouse infections were prepared by growing 50 ml cultures in LB broth (16 hours, 37°C, 200 rpm), started from a freshly grown single colony. The overnight cultures were pelleted by centrifugation and then resuspended in 5 ml of sterile PBS followed by normalization for cell number. The cultures were then promptly used to infect mice by oral gavage (200 μl per mouse). The remaining culture was used to confirm the respective inocula by plating onto LB agar. Fecal pellets were collected at Day 3, 5, 7, and 9 for each treatment group and plated on Mackonkey agar to enumerate *C. rodentium*. After 9 days, mice were sacrificed to collect colon samples. The colons were weighed, flushed with ice cold PBS to remove fibrous material. A distal 2 cm sample was weighed, homogenized in PBS, followed by serial dilution and plating onto Mackonkey agar. *Citrobacter* colonies appeared as white with pink centres after 24 hours of growth at 37°C.

### Ethics statement

The grant proposal supporting this research was reviewed and approved by the Dalhousie University Ethics Officer and the Izaak Walter Killam (IWK) research ethics committee. All animals were handled under the approval of the University Committee on Laboratory Animals under protocol #16-001, Dalhousie University which applies the guidelines of the Canadian Council on Animal Care.

## Acknowledgements

The authors acknowledge the efforts of Breanna Hargreaves, Divya Thomas, and Bruce Kuwahara with cloning steps. Alejandro Cohen provided guidance and technical support with mass spectrometry approaches and Hana James for helping with the mouse experiments. We thank members of the Thomas Lab and colleagues at Dalhousie University for critical reading of the manuscript. This work was supported by grants from the Dalhousie Medical Research Foundation (DMRF) and by equipment infrastructure from the Canadian Foundation for Innovation (CFI). The funders had no role in study design, data collection and interpretation, or the decision to submit the work for publication.

---

**Dataset S1. Basic Local Alignment Sequence Tool (BLAST) analysis of EPEC CesT compared to the non-redundant NCBI database.**

**Fig S2. Immunoblot detection of EPEC CesT, CesT variants, and effectors. (A)** The infected HeLa monolayer samples were collected 3.5 hours post infection and subjected to SDS-PAGE followed by detection with anti-CesT antibodies.

**Fig S3. Type III effector capture assay using purified recombinant histidine tagged EPEC CesT and CesT variants.** Total secreted proteins derived a Δ*sepD* strain that hypersecretes effector proteins (and is deficient for translocators) was exposed to recombinant versions of CesT as indicated (left). The imidazole elution profiles of each capture assay are indicated at right. Each fraction contains the respective CesT protein and co-eluted type III effectors as indicated. Asterisks indicate known breakdown products of Tir and NleA proteins. Note that EspC is not a type III effector and did not bind to any CesT proteins.

**Fig S4. Analysis of specific EHEC CesT substitution variants on total effector secretion profiles in an EPEC hypersecretion genetic background. (A)** Coomassie stained SDS-PAGE of total secreted proteins derived from culture supernatants of the indicated bacterial strains. **(B)** Quantitative analysis of secreted protein fractions using a Bradford assay. Absorbance readings representing total protein amounts in respective culture supernatants were divided by the optical density (OD600) of the relevant culture at time of harvest. Data are from two experiments with averages and standard deviations shown. A one-way ANOVA indicated where the data means were statistically different (P < 0.05) and a Tukey’s multiple comparison test identified confidence intervals denoted by asterisks; * = 0.045-0.32, ** = 0.064-0.34, and *** = 0.15-0.43. **(C)** Immunoblot to detect EHEC CesT or CesT variant expression levels in total cell lysates. The secreted protein profiles presented in panel A were derived from the shown lysates. Tyrosine to phenylalanine substitution CesT variants were observed to have a lower apparent molecular mass compared to EHEC CesT). The reason for this difference requires additional confirmation however other independent CesT variants have previously been observed to exhibit altered migration in protein gels This experiment was repeated three times. A representative protein gel and immunoblot are shown.

**Fig S5. Translocation dynamics for EPEC effectors that interact with CesT. (A)** EPEC strains, each containing a different chromosomally encoded effector-*blaM* gene, were subjected to kinetic translocation analysis. HeLa cells were pre-loaded with CCF2/AM and infected at an MOI of 200 with bacterial strains ‘pre-activated’ in DMEM for 4 hours. The [R] value (y-axis) is a quantitative ratio representative of the effector-TEM-1 accumulation within infected HeLa cells. Wild type EPEC carrying pCX341 (expresses unfused TEM-1) was used as a negative control for non-T3SS-specific beta-lactamase activity in a translocation competent bacterial population. Each data point represents the average from three independent experiments. **(B)** EPEC effector-*blaM* strains used in real-time translocation assays do not have significant growth differences after 4h in DMEM. Liquid overnight cultures grown in 5 mL LB+ tetracycline (10 μg/μL) were subcultured 1/200 into DMEM and incubated statically for 4h at 37°C, 5% CO_2_ to induce T3SS gene expression. OD_600 nm_ measurements were made with the Biophotometer Plus (Eppendorf) instrument. For MOI calculations based on OD_600_ (x), cell number (y) was determined with the formula y = = 746.4337(x) established from viable cell plating based on OD_600_.

**Fig S6.** Mass spectrometry analyses of CesT-FLAG LysC derived peptides. **(A)** MS1 spectrum showing a highlighted +4 charged peptide with a mass/charge ratio of 586.25 that corresponds to a specific monophosphorylated CesT peptide. **(B)** MS1 spectrum showing a highlighted +4 charged peptide with a mass/charge ratio of 566.01 that corresponds to a specific unmodified CesT peptide. **(C)** MS2 product ion spectrum for the charged peptide shown in panel A. Detected b and y fragment ions are highlighted in red and blue respectively in the table shown on the right. The enlarged inset show critical b and y ions used for site specific determination of phosphorylation on Tyr153. Mass spectrometry analyses of CesTY153F-FLAG LysC derived peptides. A) MS1 spectrum showing a highlighted +4 charged peptide with a mass/charge ratio of 582.00 that corresponds to a specific monophosphorylated CesT peptide. B) MS1 spectrum showing a highlighted +4 charged peptide with a mass/charge ratio of 562.01 that corresponds to a specific unmodified CesT peptide. C) MS2 product ion spectrum for the charged peptide shown in panel A. Detected b and y fragment ions are highlighted in red and blue respectively in the table shown on the right.

## References

1. Croxen MA, Finlay BB. 2010. Molecular mechanisms of Escherichia coli pathogenicity. Nat Rev Microbiol 8:26–38.

2. Kaper JB, Nataro JP, Mobley HL. 2004. Pathogenic Escherichia coli. Nat Rev Microbiol 2:123–140.

3. Wong AR, Pearson JS, Bright MD, Munera D, Robinson KS, Lee SF, Frankel G, Hartland EL. 2011. Enteropathogenic and enterohaemorrhagic Escherichia coli:even more subversive elements. Mol Microbiol 80:1420–1438.

4. Hazen TH, Donnenberg MS, Panchalingam S, Antonio M, Hossain A, Mandomando I, Ochieng JB, Ramamurthy T, Tamboura B, Qureshi S, Quadri F, Zaidi A, Kotloff KL, Levine MM, Barry EM, Kaper JB, Rasko DA, Nataro JP. 2016. Genomic diversity of EPEC associated with clinical presentations of differing severity. Nat Microbiol 1:15014.

5. Ritchie JM, Thorpe CM, Rogers AB, Waldor MK. 2003. Critical roles for stx2, eae, and tir in enterohemorrhagic Escherichia coli-induced diarrhea and intestinal inflammation in infant rabbits. Infect Immun 71:7129–7139.

6. Tacket CO, Sztein MB, Losonsky G, Abe A, Finlay BB, McNamara BP, Fantry GT, James SP, Nataro JP, Levine MM, Donnenberg MS. 2000. Role of EspB in experimental human enteropathogenic Escherichia coli infection. Infect Immun 68:3689–3695.

7. Marches O, Nougayrede JP, Boullier S, Mainil J, Charlier G, Raymond I, Pohl P, Boury M, De Rycke J, Milon A, Oswald E. 2000. Role of tir and intimin in the virulence of rabbit enteropathogenic Escherichia coli serotype O103:H2. Infect Immun 68:2171–2182.

8. Deng W, Puente JL, Gruenheid S, Li Y, Vallance BA, Vazquez A, Barba J, Ibarra JA, O’Donnell P, Metalnikov P, Ashman K, Lee S, Goode D, Pawson T, Finlay BB. 2004. Dissecting virulence: systematic and functional analyses of a pathogenicity island. Proc Natl Acad Sci U S A 101:3597–3602.9.

9. Pearson JS, Giogha C, Wong Fok Lung T, Hartland EL. 2016. The Genetics of Enteropathogenic Escherichia coli Virulence. Annu Rev Genet 50:493–513.

10. Pacheco AR, Sperandio V. 2012. Shiga toxin in enterohemorrhagic E.coli: regulation and novel anti-virulence strategies. Front Cell Infect Microbiol 2:81.

11. Karmali MA, Petric M, Lim C, Fleming PC, Steele BT. 1983. Escherichia coli cytotoxin, haemolytic-uraemic syndrome, and haemorrhagic colitis. Lancet 2:1299–1300.

12. Mellies JL, Barron AM, Carmona AM. 2007. Enteropathogenic and enterohemorrhagic Escherichia coli virulence gene regulation. Infect Immun 75:4199–4210.

13. Xu X, McAteer SP, Tree JJ, Shaw DJ, Wolfson EB, Beatson SA, Roe AJ, Allison LJ, Chase-Topping ME, Mahajan A, Tozzoli R, Woolhouse ME, Morabito S, Gally DL. 2012. Lysogeny with Shiga toxin 2-encoding bacteriophages represses type III secretion in enterohemorrhagic Escherichia coli. PLoS Pathog 8:e1002672.

14. Tree JJ, Wolfson EB, Wang D, Roe AJ, Gally DL. 2009. Controlling injection: regulation of type III secretion in enterohaemorrhagic Escherichia coli. Trends Microbiol 17:361–370.

15. Bustamante VH, Villalba MI, Garcia-Angulo VA, Vazquez A, Martinez LC, Jimenez R, Puente JL. 2011. PerC and GrlA independently regulate Ler expression in enteropathogenic Escherichia coli. Mol Microbiol 82:398–415.

16. Iguchi A, Thomson NR, Ogura Y, Saunders D, Ooka T, Henderson IR, Harris D, Asadulghani M, Kurokawa K, Dean P, Kenny B, Quail MA, Thurston S, Dougan G, Hayashi T, Parkhill J, Frankel G. 2009. Complete genome sequence and comparative genome analysis of enteropathogenic Escherichia coli O127:H6 strain E2348/69. J Bacteriol 191:347–354.16.

17. Perna NT, Plunkett G, 3rd, Burland V, Mau B, Glasner JD, Rose DJ, Mayhew GF, Evans PS, Gregor J, Kirkpatrick HA, Posfai G, Hackett J, Klink S, Boutin A, Shao Y, Miller L, Grotbeck EJ, Davis NW, Lim A, Dimalanta ET, Potamousis KD, Apodaca J, Anantharaman TS, Lin J, Yen G, Schwartz DC, Welch RA, Blattner FR. 2001. Genome sequence of enterohaemorrhagic Escherichia coli O157:H7. Nature 409:529–533.

18. Tobe T, Beatson SA, Taniguchi H, Abe H, Bailey CM, Fivian A, Younis R, Matthews S, Marches O, Frankel G, Hayashi T, Pallen MJ. 2006. An extensive repertoire of type III secretion effectors in Escherichia coli O157 and the role of lambdoid phages in their dissemination. Proc Natl Acad Sci U S A 103:14941–14946.

19. Hayashi T, Makino K, Ohnishi M, Kurokawa K, Ishii K, Yokoyama K, Han CG, Ohtsubo E, Nakayama K, Murata T, Tanaka M, Tobe T, Iida T, Takami H, Honda T, Sasakawa C, Ogasawara N, Yasunaga T, Kuhara S, Shiba T, Hattori M, Shinagawa H. 2001. Complete genome sequence of enterohemorrhagic Escherichia coli O157:H7 and genomic comparison with a laboratory strain K-12. DNA Res 8:11–22.

20. Shaulov L, Gershberg J, Deng W, Finlay BB, Sal-Man N. 2017. The Ruler Protein EscP of the Enteropathogenic Escherichia coli Type III Secretion System Is Involved in Calcium Sensing and Secretion Hierarchy Regulation by Interacting with the Gatekeeper Protein SepL. MBio 8.

21. Mills E, Baruch K, Charpentier X, Kobi S, Rosenshine I. 2008. Real-time analysis of effector translocation by the type III secretion system of enteropathogenic Escherichia coli. Cell Host Microbe 3:104–113.

22. Portaliou AG, Tsolis KC, Loos MS, Balabanidou V, Rayo J, Tsirigotaki A, Crepin VF, Frankel G, Kalodimos CG, Karamanou S, Economou A. 2017. Hierarchical protein targeting and secretion is controlled by an affinity switch in the type III secretion system of enteropathogenic Escherichia coli. EMBO J 36:3517–3531.

23. Thomas NA, Deng W, Baker N, Puente J, Finlay BB. 2007. Hierarchical delivery of an essential host colonization factor in enteropathogenic Escherichia coli. J Biol Chem 282:29634–29645.

24. Ramu T, Prasad ME, Connors E, Mishra A, Thomassin JL, Leblanc J, Rainey JK, Thomas NA. 2013. A Novel C-Terminal Region within the Multicargo Type III Secretion Chaperone CesT Contributes to Effector Secretion. J Bacteriol 195:740–756.

25. Brady MJ, Radhakrishnan P, Liu H, Magoun L, Murphy KC, Mukherjee J, Donohue-Rolfe A, Tzipori S, Leong JM. 2011. Enhanced Actin Pedestal Formation by Enterohemorrhagic Escherichia coli O157:H7 Adapted to the Mammalian Host. Front Microbiol 2:226.

26. Kenny B, DeVinney R, Stein M, Reinscheid DJ, Frey EA, Finlay BB. 1997. Enteropathogenic E. coli (EPEC) transfers its receptor for intimate adherence into mammalian cells. Cell 91:511–520.

27. Thomas NA, Ma I, Prasad ME, Rafuse C. 2012. Expanded roles for multicargo and class 1B effector chaperones in type III secretion. J Bacteriol 194:3767–3773.

28. Luo Y, Bertero MG, Frey EA, Pfuetzner RA, Wenk MR, Creagh L, Marcus SL, Lim D, Sicheri F, Kay C, Haynes C, Finlay BB, Strynadka NC. 2001. Structural and biochemical characterization of the type III secretion chaperones CesT and SigE. Nat Struct Biol 8:1031–1036.28.

29. Brouwers E, Ma I, Thomas NA. 2012. Dual temporal transcription activation mechanisms control cesT expression in enteropathogenic Escherichia coli. Microbiology 158:2246–2261.

30. Galan JE, Lara-Tejero M, Marlovits TC, Wagner S. 2014. Bacterial type III secretion systems: specialized nanomachines for protein delivery into target cells. Annu Rev Microbiol 68:415–438.

31. Thomas NA, Deng W, Puente JL, Frey EA, Yip CK, Strynadka NC, Finlay BB. 2005. CesT is a multi-effector chaperone and recruitment factor required for the efficient type III secretion of both LEE-and non-LEE-encoded effectors of enteropathogenic Escherichia coli. Mol Microbiol 57:1762–1779.

32. Gauthier A, Finlay BB. 2003. Translocated intimin receptor and its chaperone interact with ATPase of the type III secretion apparatus of enteropathogenic Escherichia coli. J Bacteriol 185:6747–6755.

33. Hansen AM, Chaerkady R, Sharma J, Diaz-Mejia JJ, Tyagi N, Renuse S, Jacob HK, Pinto SM, Sahasrabuddhe NA, Kim MS, Delanghe B, Srinivasan N, Emili A, Kaper JB, Pandey A. 2013. The Escherichia coli phosphotyrosine proteome relates to core pathways and virulence. PLoS Pathog 9:e1003403.

34. Abe A, de Grado M, Pfuetzner RA, Sanchez-Sanmartin C, Devinney R, Puente JL, Strynadka NC, Finlay BB. 1999. Enteropathogenic Escherichia coli translocated intimin receptor, Tir, requires a specific chaperone for stable secretion. Mol Microbiol 33:1162–1175.

35. Elliott SJ, Hutcheson SW, Dubois MS, Mellies JL, Wainwright LA, Batchelor M, Frankel G, Knutton S, Kaper JB. 1999. Identification of CesT, a chaperone for the type III secretion of Tir in enteropathogenic Escherichia coli. Mol Microbiol 33:1176–1189.

36. Deng W, Li Y, Hardwidge PR, Frey EA, Pfuetzner RA, Lee S, Gruenheid S, Strynakda NC, Puente JL, Finlay BB. 2005. Regulation of type III secretion hierarchy of translocators and effectors in attaching and effacing bacterial pathogens. Infect Immun 73:2135–2146.

37. Charpentier X, Oswald E. 2004. Identification of the secretion and translocation domain of the enteropathogenic and enterohemorrhagic Escherichia coli effector Cif, using TEM-1 beta-lactamase as a new fluorescence-based reporter. J Bacteriol 186:5486–5495.

38. Collins JW, Keeney KM, Crepin VF, Rathinam VA, Fitzgerald KA, Finlay BB, Frankel G. 2014. Citrobacter rodentium: infection, inflammation and the microbiota. Nat Rev Microbiol 12:612–623.

39. Deng W, Vallance BA, Li Y, Puente JL, Finlay BB. 2003. Citrobacter rodentium translocated intimin receptor (Tir) is an essential virulence factor needed for actin condensation, intestinal colonization and colonic hyperplasia in mice. Mol Microbiol 48:95–115.

40. Crepin VF, Collins JW, Habibzay M, Frankel G. 2016. Citrobacter rodentium mouse model of bacterial infection. Nat Protoc 11:1851–1876.

41. Buschor S, Cuenca M, Uster SS, Scharen OP, Balmer ML, Terrazos MA, Schurch CM, Hapfelmeier S. 2017. Innate immunity restricts Citrobacter rodentium A/E pathogenesis initiation to an early window of opportunity. PLoS Pathog 13:e1006476.

42. Mundy R, Petrovska L, Smollett K, Simpson N, Wilson RK, Yu J, Tu X, Rosenshine I, Clare S, Dougan G, Frankel G. 2004. Identification of a novel Citrobacter rodentium type III secreted protein, EspI, and roles of this and other secreted proteins in infection. Infect Immun 72:2288–2302.

43. Raczynski AR, Muthupalani S, Schlieper K, Fox JG, Tannenbaum SR, Schauer DB. 2012. Enteric infection with Citrobacter rodentium induces coagulative liver necrosis and hepatic inflammation prior to peak infection and colonic disease. PLoS One 7:e33099.

44. Elliott SJ, O’Connell CB, Koutsouris A, Brinkley C, Donnenberg MS, Hecht G, Kaper JB. 2002. A gene from the locus of enterocyte effacement that is required for enteropathogenic Escherichia coli to increase tight-junction permeability encodes a chaperone for EspF. Infect Immun 70:2271–2277.

45. Minamino T, Kinoshita M, Hara N, Takeuchi S, Hida A, Koya S, Glenwright H, Imada K, Aldridge PD, Namba K. 2012. Interaction of a bacterial flagellar chaperone FlgN with FlhA is required for efficient export of its cognate substrates. Mol Microbiol 83:775–788.

46. Alegria MC, Docena C, Khater L, Ramos CH, da Silva AC, Farah CS. 2004. New protein-protein interactions identified for the regulatory and structural components and substrates of the type III Secretion system of the phytopathogen Xanthomonas axonopodis Pathovar citri. J Bacteriol 186:6186–6197.

47. Yen H, Sugimoto N, Tobe T. 2015. Enteropathogenic Escherichia coli Uses NleA to Inhibit NLRP3 Inflammasome Activation. PLoS Pathog 11:e1005121.

48. Kim J, Thanabalasuriar A, Chaworth-Musters T, Fromme JC, Frey EA, Lario PI, Metalnikov P, Rizg K, Thomas NA, Lee SF, Hartland EL, Hardwidge PR, Pawson T, Strynadka NC, Finlay BB, Schekman R, Gruenheid S. 2007. The bacterial virulence factor NleA inhibits cellular protein secretion by disrupting mammalian COPII function. Cell Host Microbe 2:160–171.50.

49. Katsowich N, Elbaz N, Pal RR, Mills E, Kobi S, Kahan T, Rosenshine I. 2017. Host cell attachment elicits posttranscriptional regulation in infecting enteropathogenic bacteria. Science 355:735–739.

50. Page AL, Parsot C. 2002. Chaperones of the type III secretion pathway: jacks of all trades. Mol Microbiol 46:1–11.

51. Wattiau P, Bernier B, Deslee P, Michiels T, Cornelis GR. 1994. Individual chaperones required for Yop secretion by Yersinia. Proc Natl Acad Sci U S A 91:10493–10497.

52. Stebbins CE, Galan JE. 2001. Maintenance of an unfolded polypeptide by a cognate chaperone in bacterial type III secretion. Nature 414:77–81.

53. Lara-Tejero M, Kato J, Wagner S, Liu X, Galan JE. 2011. A sorting platform54. determines the order of protein secretion in bacterial type III systems. Science 331:1188–1191.

54. Laemmli UK. 1970. Cleavage of structural proteins during the assembly of the head of bacteriophage T4. Nature 227:680–685.

55. Gauthier A, de Grado M, Finlay BB. 2000. Mechanical fractionation reveals structural requirements for enteropathogenic Escherichia coli Tir insertion into host membranes. Infect Immun 68:4344–4348.

56. Thomassin JL, He X, Thomas NA. 2011. Role of EscU auto-cleavage in promoting type III effector translocation into host cells by enteropathogenic Escherichia coli. BMC Microbiol 11:205.

